# Retrieval and Savings of Contextual Fear Memories Across an Extended Retention Interval in Juvenile and Adult Male and Female Rats

**DOI:** 10.1101/2022.10.28.514119

**Authors:** Natalie Odynocki, Zerah Isaacs, Andrew M. Poulos

## Abstract

Adult rodents exhibit an exceptional ability to retrieve context fear memories across lengthy retention intervals. In contrast, these same memories established in juvenile rodents are susceptible to significant forgetting. While much of this original work has focused on male rodents, emerging research indicates that the mechanisms of context fear conditioning in females may be sexually dimorphic. If so, these female memories may exhibit a unique mnemonic profile from male rats that may be probed using single-trial conditioning, extended retention intervals, and savings procedures. The current study aimed to examine the longevity of contextual fear memories established in 24-day-old juvenile and 90-day-old Long-Evans male and female rats. Testing at the 1-day retention interval, adults exhibited greater conditioning than juvenile males, while in females, conditioning did not differ between juvenile and adult rats. In adults, males displayed greater conditioning than females, while in juveniles, males and females reached similar conditioning levels. At the 60-day retention interval, adult sex differences were maintained; however, juvenile rats failed to retrieve this remote contextual fear memory. Next, we examined whether a savings trial and test procedure could recover these remotely established juvenile memories in both males and females. Following a 60-day retention test, the now adult rats were presented with an additional context-shock pairing to assess the level of saving in both male and female rats. While this procedure produced greater conditioning in males than females, the relative amounts of the savings of this early life memory were similar in males and females. Lastly, the recovery of this memory exhibited significant generalization relative to previously unshocked juveniles and was similar in males and females. The results of these experiments indicate that adult sex differences in contextual fear memory are maintained across an extended retention interval, while in juveniles, there were no significant sex differences. In addition, male and female rats failed to retrieve an initial juvenile memory at an extended retention interval. However, these memories could be recovered with a single reminder of the original juvenile experience.

## Introduction

Successful retrieval of memories acquired in juvenile and adult mammals can markedly differ after prolonged periods. Adult rodents demonstrate an exceptional ability to retrieve context fear memories across lengthy retention intervals extending well beyond a year (Gale et al., 2004). In contrast, these same memories established in juvenile rodents are susceptible to rapid forgetting yet can be reinstated with reminder treatments (Rudy & Morledge, 1994; Kim et al., 2009; Akers et al., 2014; Alberini & Travaglia, 2017). While much of this original work has focused on males or a mixture of male and female rodents, a growing body of research indicates that the neuro-cognitive mechanisms of context fear conditioning in females may markedly differ (Keiser et al., 2017; Colón et., 2018; Colón & Poulos, 2020). If so, these female memories may exhibit a unique profile of memory recovery from male rats than can be uncovered by examining the long-term retrieval and savings of juvenile and adult context fear memories.

Females have been under-represented as subjects in human and animal studies, particularly in the neurosciences (Beery & Zucker, 2011; Mamlouk, Dorris, Barrett, & Meitzen, 2020; Woitowich, Beery, & Woodruff, 2020; Zucker & Beery, 2010). Given that women are 2-3 fold more likely to develop fear-related disorders (Kessler et al., 2005, 2010), it is crucial to begin elucidate the processes and mechanisms that underlie this disparity by including females in the study of fear conditioning.

In recent years, the inclusion of female subjects has led to identifying adult sex differences in context fear conditioning and extinction in rats and mice (Maren et al., 1994; Valero et al., 2019; Colón et al., 2020; Binette et al., 2022). Comparative studies of adult male and female animals have found that males exhibit greater contextual fear conditioning than females (Maren et al., 1994; Wiltgen et al., 2001; Gresack et al., 2009; Colón et al., 2018; Colón et al., 2020). We recently proposed that these adult differences are due to different context-spatial processing demands exerted on the hippocampus (Colón & Poulos, 2020). Specifically, increasing contextual encoding in males correspondingly engages a greater number of CA1 neurons, while in females, this did not significantly alter the recruitment of CA1 neurons. Interestingly, animals with lesions of the hippocampus that include CA1 can acquire contextual fear memories; however, they exhibit deficits in retrieval that become more severe with longer retention intervals (Zelikowsky et al., 201X). In intact adult females, this natural reduction of hippocampal engagement during acquisition and retrieval may promote a decline in the retrieval of contextual fear memories that increases with the length of the acquisition-to-retention test interval that is not present in males.

In rats, the development of contextual fear conditioning typically emerges at approximately 21 to 24 days postnatal (P) (Rudy, 1993; Burman et al., 2009; Santarelli et al., 2018). However, during this juvenile period, learning progresses more slowly than in adults (Colón et al., 2018) and is susceptible to considerable forgetting at longer retention intervals (Rudy & Morledge, 1994; Robinson-Drummer & Stanton, 2015). It has been proposed that adult-like levels of conditioning and memory retrieval are not attained until the hippocampus fully develops, yet these early memories remain accessible with sufficient reminder cues (Travaglia et al., 2016). We recently found evidence that hippocampal CA1 of juvenile males is not significantly engaged in retrieving contextual fear memories as in adult males (Santarelli et al., 2018). In a separate study, we found that juvenile females can display greater context fear conditioning than age-matched males and litter-matched adult females. This learning quality in juvenile females, not evident in males, may promote a greater endurance of contextual fear memories across extended retention intervals.

The present experiment examines the longevity of contextual fear memories in juvenile and adult male and female rats. If learning is indeed enhanced in juvenile females, testing at a long retention interval should alleviate forgetting evident in juvenile males. Conversely, if contextual fear learning in adult females does not significantly engage the hippocampus as it does in adult males, testing at a remote retention interval, which requires the hippocampus, ought to result in poorer retrieval in females, not evident in males. Finally, if forgetting is due to a retrieval failure, in groups that exhibit forgetting, savings of these memories should be evident with a sufficient reminder trial.

In the following experiment, we used a single-trial context fear conditioning procedure in litter- and age-matched P24 juvenile and P90 adult male and female Long-Evans rats. Memory retention was tested at a 1 day or extended retention interval of 60 days. To further probe forgetting, we employed a savings trial, which consisted of an additional footshock in the original context, followed by a savings test of context fear in the same and novel context.

## Methods

### Subjects and Housing

A total of 178 Long Evans rats were included in experiment 1, and 45 were included in experiment 2. Rats were bred in-house within the Life Sciences Research Building vivarium at the University at Albany, SUNY. Rats were kept on a 14:10 light/dark cycle with ad libitum access to food and water. Litters were weaned on postnatal day (P) 21 and cohoused with same-sex littermates. Rats were transported to a holding room separate from the conditioning room, where they were habituated to handling for 5 minutes each day for three consecutive days leading up to acquisition training. All experiments were carried out during the light phase. Animal care and experiments were conducted in adherence to protocols approved by the Institutional Animal Care and Use Committee (IACUC) at the University at Albany.

### Apparatus

Context A consisted of four sound-attenuating chambers measuring 30.5×24 1×21 cm (Med Associates Inc.). The floor of each chamber consisted of either 36 stainless steel rods (29.3×26×6.1 cm) with bars 7.9 mm in diameter spaced 1.8 cm apart for P24 subjects or 19 stainless steel rods (29.3× 26×6.1 cm) with bars 16 mm in diameter spaced 2.8 cm apart for P90 subjects wired to a shock generator. Before conditioning or context retrieval, drop pans were scented with 50% Simple Green Solution and inserted under the grid floors to provide an olfactory cue. White fluorescent light placed above the testing chamber and a 60-dB ventilation fan located in the back of the chamber provided ambient light and noise, respectively. The walls of the context were plain white except for the front, which is a transparent door. This context was housed within a chamber to minimize external sound and light sources.

Context B consisted of the same chambers and dimensions as context A but was designed to be distinct from context A. A smooth acrylic white floor insert was placed over stainless-steel grid floors. A black plastic A-frame insert spanned from the grid floor to the ceiling to give the context a triangular shape. Before and after each testing session, all chambers were cleaned with 25% isopropyl alcohol. Drop pans were scented with 1% acetic acid and inserted under the grid floors. Red fluorescent light illuminated the room outside the chambers, and ventilation fans at the back of each chamber were turned off to eliminate background noise.

## Procedures

### Experiment 1

On acquisition day, P24 and P90 male and female rats were transported to the same holding room where they were handled the three previous days. Rats were habituated in the holding room after transport for at least 15 minutes before being placed in the conditioning context for the first time. Rats were then brought to the conditioning room and placed in conditioning context A, where they remained for 4 minutes and 2 seconds. After a 3-minute baseline period, a 1.25 mA (2 sec) footshock was administered through a grid floor. Rats remained in the chamber for an additional minute before being returned to their home-cage. In a control group, rats were exposed to the context but received no footshock. At 1 or 60 days post-acquisition, rats were returned to the training context for a 4-minute recent or remote retrieval test, respectively.

### Experiment 2

On acquisition day, P24 male and female rats were transported to the holding room and conditioned in a manner identical to experiment 1, including a non-shock control group. Sixty days post-acquisition, rats were returned to context A for a 4-minute retrieval test. At the end of 4 minutes, all rats, including those in the non-shock control group during acquisition, received a 1.25mA (2 sec) footshock and remained in the context for an additional minute as a savings trial. Rats shocked at both acquisition and the end of the 60-day retrieval test will be referred to as the shk-shk group. Rats that were not shocked at acquisition but later shocked at the end of the 60-day retrieval will be referred to as the ns-shk group. Rats were then returned to their home cages and transported to the vivarium. Twenty-four hours later, rats were returned to context A for a 4-minute savings test and then returned to the vivarium. Twenty-four hours after the savings test, rats were placed in context B for a 4-minute test for the context specificity of savings.

### Freezing

Behavior during retrieval tests was video recorded using a near-infrared camera mounted to the door of the sound attenuating box. Freezing was defined as the cessation of all movement except those related to respiration. Freezing during recent and remote retrieval was manually measured by an experimenter blind to the shock condition of the animal. The experimenter used a stopwatch to record the number of seconds the rat spent freezing. The number of seconds of freezing was then converted into the percent time spent freezing by dividing the number of seconds spent freezing by the total number of seconds tested in the chamber during retrieval (240 sec).

## Statistical Analysis

### Experiment 1

A 4-way ANOVA was run in SPSS for this 2 × 2 × 2 × 2 factorial design to detect any combination of interactions between age (P24 vs. P90), sex, retention interval (1-day vs. 60 days), and shock condition (shock vs. no shock). If there were any significant interactions, we tested for simple main effects.

### Experiment 2

A 2-way ANOVA was run in SPSS to detect interactions between shock condition and sex or 60-day retrieval, savings test, and generalization test. An examination of simple main effects followed significant interactions.

## Results

### Experiment 1

A 3-way interaction between age, sex, and shock was observed F (1,15) =6.144, p=.014, along with two-way interactions between retrieval by shock (F (1,15) =5.631, p=.019) and retrieval by age (F (1,15) =11.271, p=.001). Simple main effects revealed that irrespective of sex, shocked P24 animals tested at the recent retention interval exhibited greater free zing than non-shocked animals (F (2,159) =10.57, p=.000). Interestingly, there was no significant difference at the remote interval, as shocked animals froze at an equal rate to non-shocked animals, suggesting forgetting. This was further supported in P24 shocked animals, exhibiting greater freezing at the recent interval than at the remote (F (2, 159)=37.21, p=.000).

In P90 shocked animals, males displayed higher levels of freezing compared to females at both recent (F (1,159) = 31.2 6, p=.000) and remote (F (1,159) =35.02, p=.000) intervals. Across ages, P90 shocked males showed greater freezing than P24 shocked males at both the recent (F (1,1 59) =50.30, p= 000) and remote (F (1,159) =122.98, p=.000) retention intervals. Conditioned P90 females froze more than P24 females at the remote retention interval (F (1,159) =27.84), p=.000).

### Experiment 2

No significant interactions between sex or shock condition were observed at the 60-day test of retrieval, savings, and generalization. For the savings test, an effect of sex was identified (F (1,42) =15 .024, p=.000), wherein males showed greater evidence of savings than females. There was also a main effect of shock (F (1,42) =20.601, p=.000) in which animals shocked initially during acquisition (shk-shk) had a greater propensity to freeze during the savings test than animals that were in the non-shock condition during acquisition (ns-shk). In the absence of an interaction, this suggests no statistical difference in the proportion of savings between the shk-shk and ns-shk groups. To test the specificity of savings, freezing was assessed in Context B, where a main effect of shock (F (1,42) =15.674, p=.000) was identified, in which animals that were shocked during acquisition (shk-shk) generalized more than animals not shocked at acquisition (ns-shk). In addition, no main effects of sex were identified.

## Discussion

The present study examined the long-term retention of contextual fear memory in juvenile and adult female rats compared to age-matched male rats. We believe these results facilitate our understanding of the developmental emergence of sex differences in the long-term retrieval of contextual fear memories. The current results replicate previous findings of adult sex differences in context fear learning and provide novel insights into long-term maintenance and retrieval of these memories in juvenile male and female rats.

In experiment 1, we tested our initial hypotheses that adult female rats tested at a remote retention interval would exhibit poorer retrieval, which would not be evident in adult males, and that juvenile female rats tested at a remote retention interval would demonstrate superior retrieval than juvenile males. Contrary to these hypotheses, we found that male and female adult animals exhibited a sustained level of conditioned fear across a 1- to 60-day retention interval, with males demonstrating consistently higher levels of fear than females. Furthermore, we found that irrespective of sex, conditioned juveniles froze at significantly lower levels than adults. By the 60-day retention interval, conditioned juveniles froze at levels equal to that of non-conditioned animals, indicating that juveniles failed to retrieve the initial contextual fear memory.

Adult sex differences in context fear conditioning were evident at the one and 60-day retention intervals. In particular, the decreased context fear learning in females at the recent test was not only sustained after two months but also exhibited no evidence of forgetting as previously described in male rodents. These results can be attributed to male and female differences in hippocampal-dependent spatial learning (Yagi & Galea, 2019). Recently, we proposed that in adult males that hippocampal CA1 neurons may more readily process contextual-spatial information than in adult females (Colón et al., 2020). In this study, increasing context exposure time increased the number of Fos-immunoreactive (ir) neurons in the hippocampal CA1 of male but not female mice at both conditioning and testing. These sex differences are further evident using molecular and electrophysiological measures of synaptic plasticity. Kudo et al. (2004) demonstrated in males that context fear conditioning increased the phosphorylation of CREB in hippocampal CA1 neurons that were not evident in females. At the same time, Maren et al. (1994), using *in-vivo* electrophysiological stimulation and recordings from the hippocampus, demonstrated that long-term potentiation (LTP) is reduced in females relative to males. In addition, these differences may be related to levels of estrogens at the time of conditioning and testing that naturally fluctuate at different phases of the estrous cycle in females. However, it should be noted that several studies have found that the phase of the estrous cycle does not readily affect the acquisition or retrieval of contextual fear memories (Blume et al., 2017; Cossio, Carreira, Vásquez, & Britton, 2016; Keiser et al., 2017; Maeng & Milad, 2015; R, Machado, et al., 2019).

In juvenile and adult rats, we observed a sex-specific development of context fear conditioning. In male rats, this was particularly evident in the 24-hour test of context fear, where conditioned responding in adults was greater than in juvenile rats, a result consistent with prior studies (Colon et al., 2018; Santarelli et al., 2018). We recently proposed that this developmental difference may reflect a unique contribution of the prefrontal cortical regions in juvenile contextual fear learning. In support of this view, we previously demonstrated in male rats that c-Fos within the prelimbic area exhibit a retrieval-dependent expression in juvenile rats, which is not evident in peripubertal and adult rats (Santarelli et al., 2018). Consistent with this, inactivation of the prelimbic area in adult male rats fails to disrupt context fear learning (Zelikowsky et al., 2013), while similar inactivation in juvenile rats disrupts the acquisition of contextual fear associations in both male and female rats (Heroux et al., 2017; Santos et al., 2019). Whether the contributions of the prelimbic area to context fear conditioning is specific to juvenile-age rats remains to be fully tested.

Both male and female juvenile rats exhibited a precipitously decline in freezing between the 1 (male: 13.7%, female: 20 .4%) and 60-day (male: 0.7%, female: 1.1%) retention intervals. This amnesia of early but not adult life experiences has been demonstrated in many forms of learning and across several species of animals, including humans (Campbell et al., 1966; Spear & Parsons, 1976; Haroutunian & Riccio, 1977; Davis & Rovee-Collier, 1983; Akers et al., 2012). In recent years, infantile or childhood amnesia has been attributed to the protracted maturation of the hippocampus (Akers et al., 2014; Alberini & Travaglia, 2017). During early juvenility, rates of postnatal hippocampal neurogenesis peak before attenuating to adult levels (Kuhn, Dickinson-Anson, & Gage, 1996; Seki & Arai, 1995). Experimental manipulations that reduce the rate of hippocampal neurogenesis in young mice improve the long-term retrieval of contextual fear memories, while in adult mice, procedures that enhance the rate of neurogenesis promote forgetting (Akers et al., 2014). Akers et al. (2014) have proposed that the incorporation of a larger number of newly born neurons remodels hippocampal circuits leading to an overwriting of memory traces, which promotes a decrease in memory accessibility. Conversely, Travaglia et al. (2016) have proposed that during infancy, prolonged activation of BDNF pathways and mGluR5-dependent switch of NMDA receptors subunit expression from 2B to A within the dorsal hippocampus promotes forgetting in a context-dependent version of the inhibitory avoidance that can be recovered.

In our second experiment, we addressed the hypothesis that although juvenile contextual fear memory does not appear to be retrieved at the 60-day retention interval, this early life memory is indeed intact and, to some extent, recoverable. Using a savings trial and test procedure (Ebbinghaus, 1885; Poulos et al., 2009), we found evidence in support of retrieval failure accounts of forgetting, with savings evident in both adult male and female rats initially conditioned during juvenility.

Consistent with previous accounts of forgetting and as demonstrated here in both male and female rats, these early memories are recoverable and highlight a failure to retrieve rather than a loss of the original memory (Alberini & Travaglia, 2017; Anderson & Riccio, 2005; Guskjolen et al., 2018; Jasnow et al., 2012; Li et al., 2014; Rudy & Morledge, 1994). In the current study, a savings trial could alleviate the initial failure to retrieve a juvenile contextual fear memory across an extended interval. In experiment 2, after a 60-day retention interval, all animals, previously conditioned or not during juvenility, were subjected to a single shock in the original juvenile context and given a context test 24 hours. The juvenile-conditioned group exhibited greater freezing than the group not previously conditioned, providing evidence for the “savings” of the original juvenile memory. The extent of greater freezing was proportional in male and female subjects suggesting that this juvenile memory or its storage was similar in male and female rats. Lastly, to assess the context-specificity of juvenile recovered memory, subsequent testing in a novel context, both males and females exhibited considerable freezing over juvenile naïve controls. This effect in juvenile trained animals may highlight that remotely acquired memory, over those more recently acquired, are more prone to another form of retrieval failure, notably evident in adults, context generalization (Wiltgen & Silva, 2007; Biedenkapp & Rudy, 2007).

This study explored memory retrieval in male and female subjects at two developmental periods: juvenility and adulthood. In adulthood, sex differences in context fear conditioning were not only evident after a short-retention interval but remained so following a two-month retention interval. While no sex differences were evident in juvenile-conditioned animals, males and females initially failed to retrieve these memories after a two-month test interval. However, consistent with previous studies, these juvenile memories, irrespective of sex, appear to remain accessible well into adulthood. The current results suggest that, unlike in adults, juvenile fear-based contextual memories, while not readily retrieved after long periods, are latently stored and can remerge with similar experiences later in life.

**Figure 1:**
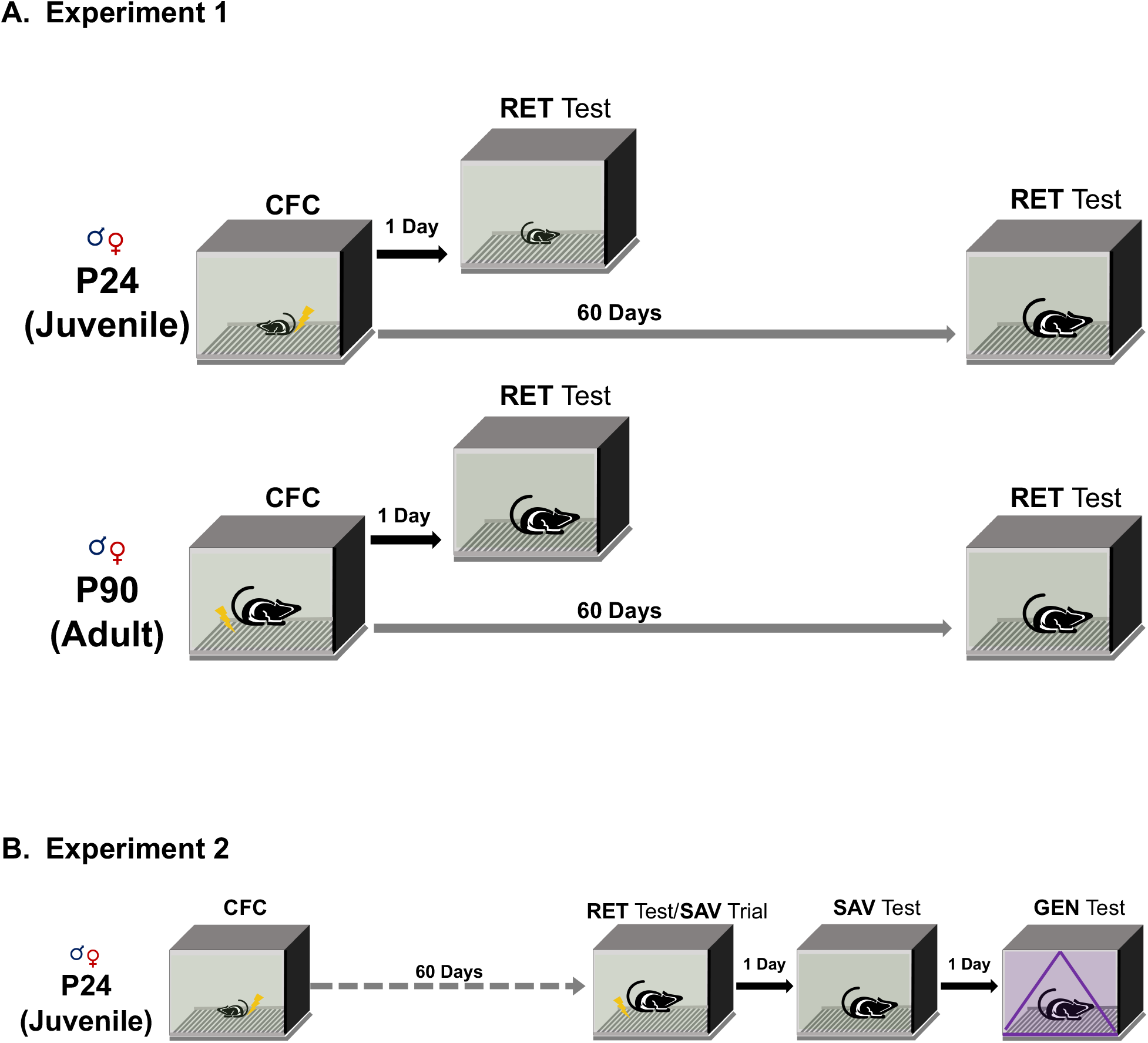
***(A)*** In experiment 1, P24 or P90 male and female Long Evans rats were conditioned in context A and returned 1 or 60 days later for recent or remote retrieval testing. ***(B)*** In experiment 2, P24 male and female rats were conditioned in context A and returned 60 days later for retrieval testing. One minute and 2 seconds before being removed from context A, all rats were administered a 1.25mA reminder footshock. Twenty-four hours later, rats were returned to context A for a savings test. Twenty-four hours after the savings test, rats were placed in context B for a context specificity test.

**Figure 2:**
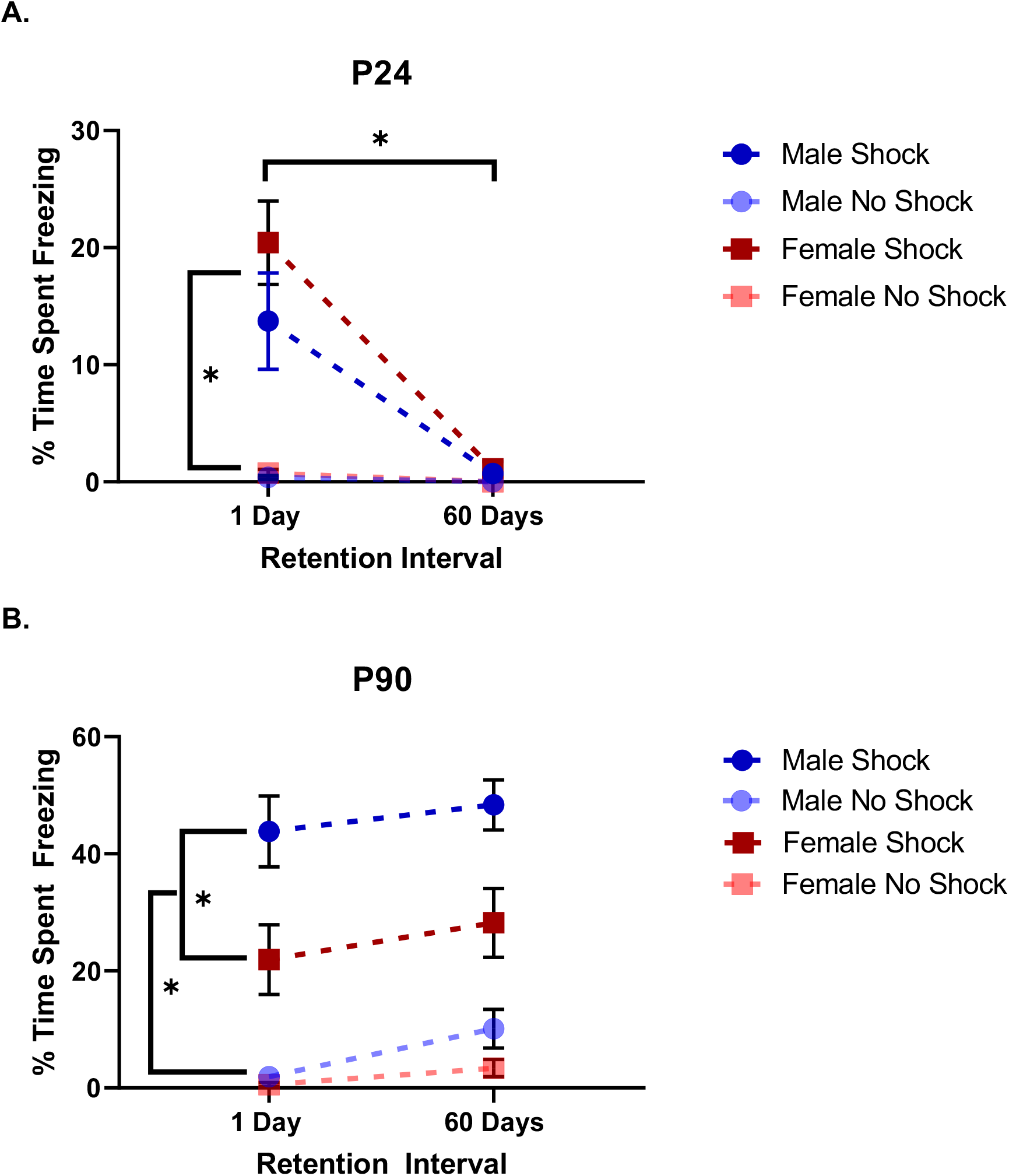
Context fear retrieval at one day and 60 days in male and female rats fear conditioned at P24 ***(A)*** and P90 ***(B)***, respectively. A 3-way ANOVA interaction between age, sex, and shock was observed F(1,15)=6.144, p=.014, along with two-way interactions between retrieval and shock (F(1,15)=5.631, p=.019) and retrieval by age (F(1,15)=11.271, p=.001). ***(A)*** Simple main effects revealed that irrespective of sex, shocked P24 animals tested at the recent retention interval exhibited greater freezing than non-shocked animals (F(2,159)=10.57, p=.000). No statistically significant differences were observed in any group at the remote interval, as shocked animals froze at a similar level to non-shocked animals, suggesting forgetting. Shocked P24 animals further support this, irrespective of sex, exhibiting a greater fear response at the recent interval than at the remote (F(2, 159)=37.21, p=.000). ***(B)*** In shocked P90 animals, males displayed greater levels of conditioned fear compared to females at both recent (F(1,159)=31.26, p=.000) and remote (F(1,159)=35.02, p=.000) intervals. Across ages, shocked P90 males showed a greater fear response than shocked P24 males at both the recent (F(1,159)=50.30, p=.000) and remote (F(1,159)=122.98, p=.000) retention intervals. P90 females only outperformed P2 4 females at the remote retention interval (F(1,159)=27.84), p=.000).

**Figure 3:**
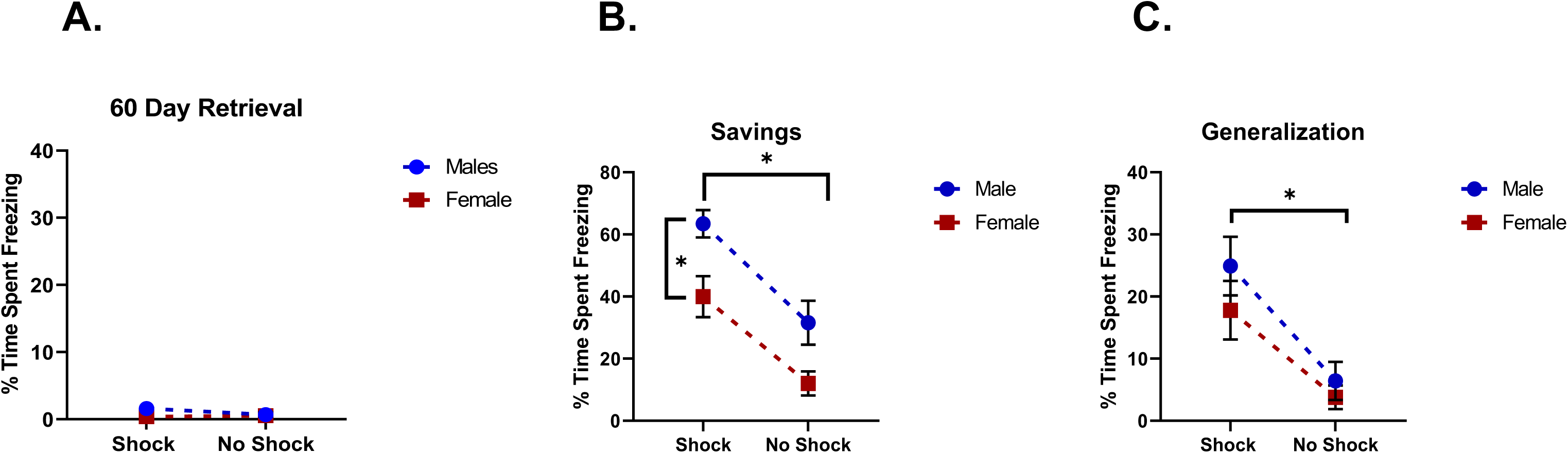
Experiment 2. ***(A)*** Animals previously shocked showed low freezing levels similar to non-shock animals, suggesting forgetting across the 60-day retention interval. ***(B)*** Savings in adult male and female rats conditioned at P24. Rats were given a single reminder shock at the end of the 60-retrieval test and returned to context A 24 hours later for a savings test. A main effect of sex (F(1,42)=15.024, p=.000) was identified, wherein males showed greater freezing than females. There was also a main effect of shock (F(1,42)=20.601, p=.000) in which animals shocked initially during acquisition (shk-shk) had a greater propensity to freeze during the savings test than animals who were in the non-shock condition during acquisition (ns-shk). The absence of an interaction suggests that savings across groups were proportionate. ***(C)*** Rats were placed in context B to test the specificity of context fear 24 hours after a savings test. A main effect of shock (F(1,42)=15.674, p=.000) was observed in which animals shocked during acquisition (shk-shk) generalized more than animals not shocked at acquisition (ns-shk), but no main effects of sex were found.

## Notes

### Competing Interest Statement

The authors have declared no competing interest.

